# A niche-specific role for ArlRS enables *Staphylococcus aureus* growth within Kupffer cells

**DOI:** 10.64898/2026.10.07.757369

**Authors:** Josefien W. Hommes, Lindsey M. Orthner, Fardau van der Wal, Ronald S. Flannagan, Raymond Shim, Luisa Samaniego Barrón, Mai Dao, Iñigo Lasa, Alexander R. Horswill, David E. Heinrichs, Bas G. J. Surewaard

## Abstract

*Staphylococcus aureus* bloodstream infections cause severe morbidity and mortality despite antibiotic therapy. We previously identified Kupffer cells (KCs) as the primary hepatic macrophages that rapidly clear circulating *S. aureus*, yet a subset of bacteria survives intracellularly and seeds systemic infection. Here, using spinning-disk intravital and super-resolution microscopy, we show that bacterial two-component system (TCS) signaling is required for intracellular replication within KCs *in vivo*. A global mutant lacking all non-essential TCSs failed to form intracellular microcolonies, and targeted analysis identified the ArlRS system as specifically required for KC replication. Δ*arlRS* bacteria remained as single non-replicating organisms within KCs, exhibited reduced dissemination and kidney abscess formation, and showed attenuated virulence with improved host survival. Complementation with *arlRS* restored the wildtype phenotype. Notably, the intracellular replication defect was not observed in primary or cultured macrophages, indicating that ArlRS-dependent replication is highly niche-specific and not reproduced by conventional *in vitro* models. Furthermore, restricting the KC reservoir with vancomycin treatment after infection further reduced renal infection by Δ*arlRS* bacteria. These findings identify ArlRS as a critical regulator of intracellular replication within KCs and establish the KC intracellular reservoir as a key driver of *S. aureus* pathogenesis and a potential therapeutic target during bloodstream infection.

## Introduction

*S. aureus* bloodstream infection is one of the most lethal bacterial infections encountered in modern medicine and remains a major challenge in the era of antimicrobial resistance. Recent studies estimate that *S. aureus* bacteremia causes hundreds of thousands of deaths annually worldwide, with case-fatality rates of approximately 15–30% despite antibiotic therapy^1,2^. In Canada, bloodstream infections caused by methicillin-resistant *S. aureus* (MRSA) were associated with approximately 20% mortality between 2018 and 2022^3^. Beyond acute mortality, *S. aureus* bacteremia is frequently complicated by metastatic infection, sepsis, and relapse, implying that the pathogen can persist in protected host niches^1^. Although *S. aureus* was classically viewed as an extracellular pathogen, we previously demonstrated that, after entering the bloodstream, *S. aureus* can establish an intracellular reservoir within Kupffer cells (KCs), the resident macrophages of the liver^4^. Within this niche, approximately 10% of bacteria survive, replicate, and eventually re-enter the circulation, driving systemic dissemination and mortality.

Intracellular persistence likely provides multiple advantages to *S. aureus*, including evasion from neutrophil-mediated killing, shielding from extracellular immune responses, and reduced accessibility to antibiotics^5,6^. Multiple studies have shown that *S. aureus* can survive within neutrophils and macrophages, where intracellular localization may facilitate dissemination and protection from antimicrobial therapy^7^. Intracellular localization has also been linked to recurrent and chronic infection, as many antibiotics exhibit poor intracellular activity despite effective extracellular killing^7,8^. Consistent with this concept, we previously demonstrated that vancomycin, the current standard-of-care therapy for MRSA bacteremia, poorly penetrates infected Kupffer cells (KCs), allowing intracellular bacteria to survive despite adequate therapy^4^.

In addition to our previous findings *in vivo*, numerous *in vitro* studies have demonstrated that *S. aureus* can survive within endothelial cells, epithelial cells, osteoclasts, neutrophils, and macrophages, supporting the idea that intracellular survival is an important immune evasion strategy during invasive disease^9–16^. To facilitate intracellular persistence, *S. aureus* employs multiple evasion strategies to withstand the hostile intracellular environment, including resistance to oxidative stress, antimicrobial peptides, nutrient limitation, and phagolysosomal killing^17–19^. However, the bacterial pathways that specifically enable *S. aureus* to establish and maintain the intracellular KC reservoir remain poorly understood.

Bacterial two-component systems (TCSs) are major sensory and regulatory pathways that enable bacteria to rapidly adapt to environmental stress. *S. aureus* encodes 16 TCSs, many of which regulate virulence, metabolism, antibiotic resistance, and stress adaptation^20,21^. Several TCSs have also been implicated in intracellular survival, including GraRS in macrophages, SaeRS and Agr in neutrophils and macrophages, and SrrAB during oxidative and nitrosative stress^22–26^. Whether specific TCSs regulate intracellular survival within tissue-resident macrophages during bloodstream infection *in vivo*, however, remains unknown.

Here, using spinning-disk intravital microscopy (SD-IVM), super-resolution imaging, and targeted bacterial genetics, we identify the ArlRS TCS as a critical determinant of intracellular *S. aureus* replication within KCs. Loss of ArlRS abolished bacterial microcolony formation within the liver, reduced systemic dissemination, and markedly improved host survival during bloodstream infection. Strikingly, this phenotype was not reproduced in conventional *in vitro* macrophage infection models, highlighting the highly specialized nature of the KC intracellular niche. We adapted a tissue-clearing protocol to visualize the entire kidney together with the presence of any formed bacterial communities three-dimensionally in high resolution. This novel approach of abscess visualization showed a stark decrease of staphylococcal presence in kidneys upon the loss of ArlRS. Together, these findings identify ArlRS as a previously unrecognized regulator of intracellular persistence *in vivo* and establish the intracellular KC reservoir as a central driver of *S. aureus* pathogenesis during bacteremia.

## Results

### Kupffer cells rapidly capture circulating *S. aureus* but permit intracellular persistence

KCs represent the largest population of tissue-resident macrophages in the body and account for ∼80% of all resident macrophages^27,28^. Strategically positioned within the liver sinusoids, KCs continuously survey the bloodstream and function as a critical intravascular barrier responsible for rapid clearance of invading intravascular bacteria^4,29,30^. Previous studies have shown that KCs are highly specialized for the capture of bloodborne microbes through CRIg-dependent or antibody-dependent interactions under shear flow conditions^31^. Using SD-IVM, we observed that KCs actively extend membrane protrusions into the sinusoidal lumen to rapidly catch circulating *S. aureus* MW2 almost immediately following intravenous (i.v.) infection (Fig. 1a, Video 1). Bacterial catching occurred very rapidly, as virtually no circulating bacteria could be detected 10 minutes after infection, and nearly all bacteria were localized intracellularly within KCs. To directly assess the importance of KCs during bloodstream infection with *S. aureus*, we selectively depleted KCs in Clec4f-Cre-tdTomato-iDTR mice by diphtheria toxin administration before infection^32^, which was confirmed with SD-IVM (Supplemental Fig. 1a). KC depletion resulted in a marked reduction in hepatic bacterial catching compared to saline-treated controls (Fig. 1b, Video 2), confirming that KCs are the dominant cell population responsible for removing *S. aureus* from the circulation. Consistent with previous reports demonstrating an essential role for KCs in protection against bacteremia^29,33^, loss of these macrophages significantly increased mortality following bloodstream infection (Fig. 1c). Although most bacteria are efficiently eliminated following KC uptake, our previous work demonstrated that a subset of intracellular *S. aureus* survives within these macrophages and establishes a protected intracellular reservoir. Indeed, SD-IVM imaging revealed the formation of large grape-like intracellular bacterial microcolonies within individual KCs 10 hours after infection (Fig. 1d). Intracellular localization within KCs protects bacteria from recruited immune cells and limits accessibility to antibiotic therapy, thereby providing a potential explanation for relapsing and metastatic disease during bacteremia^4^. However, the bacterial pathways that enable *S. aureus* to survive and replicate within this highly hostile intracellular environment remain unknown.

**Figure 1.**
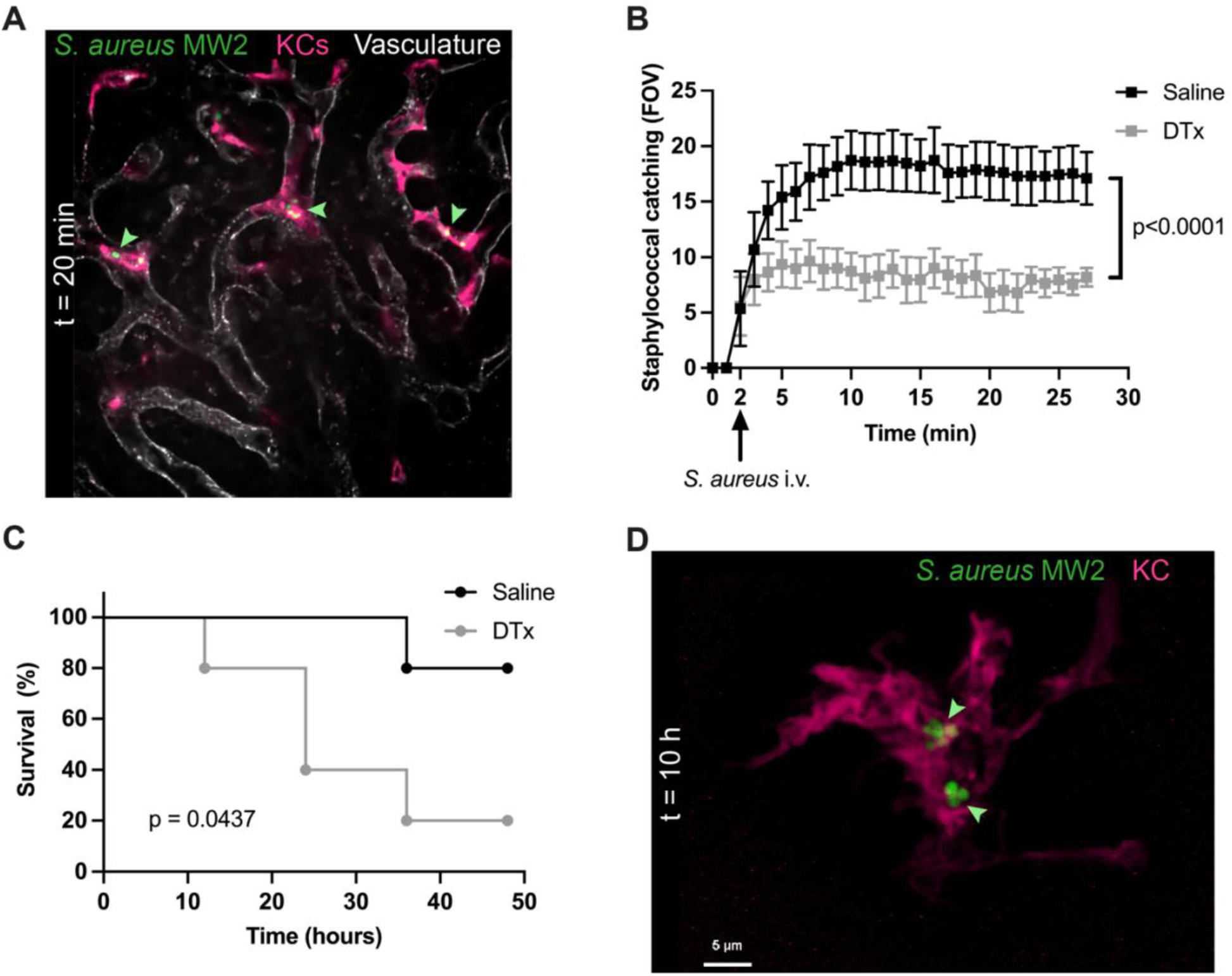
Kupffer cells are required for *S. aureus* capture during bloodstream infection. (a) Representative SD-IVM image 20 minutes after intravenous *S. aureus* infection. Image corresponds to **video S1**. Green arrows indicate *S. aureus* internalized by Kupffer cells. Vasculature (CD31; white), Kupffer cells (TIM-4; magenta), *S. aureus* MW2 (GFP; green). (b) Quantification of bacterial catching in saline-treated (black) and diphtheria toxin-treated (grey) Clec4f-Cre-TdTomato^+/-^/iDTR^+/-^ mice (n = 5 for saline and n = 6 for DTx). Welch’s t test, mean +/-SEM. Data corresponds to **video S2** (c) Survival of saline-treated (black) versus diphtheria toxin-treated (grey) Clec4f-Cre-TdTomato^+/-^/iDTR^+/-^ mice following intravenous infection (5×10^6^ CFU *S. aureus* MW2-GFP) (n = 5). Kaplan-Meier survival analysis with Mantel-Cox test. (d) Confocal microscopy whole-mount liver image 10 hours after *S. aureus* bloodstream infection showing intracellular bacterial microcolonies (green arrows). Kupffer cells (F4/80; magenta), *S. aureus* MW2 (GFP; green). Scale bar 5 µm.

### *Staphylococcus aureus* TCSs are required for intracellular survival and replication

Bacteria use TCSs to sense and adapt to environmental stimuli through rapid, coordinated changes in gene expression. As the intracellular KC niche exposes bacteria to multiple stresses, including antimicrobial proteins, nutrient limitation, acidic components and oxidative stress, we hypothesized that TCS signaling is required for *S. aureus* adaptation to the hostile intracellular KC environment. To test this, we used the clinically relevant community-associated MRSA (CA-MRSA) strain MW2, isolated from a patient with bacteraemia, and a corresponding mutant lacking all 15 non-essential two-component systems (ΔXV)^34^. SD-IVM showed significantly less bacterial microcolony formation in the livers of mice infected with ΔXV compared to wildtype *S. aureus* MW2 (Fig. 2a, b). Not only was there a reduction in microcolonies, but the average size of ΔXV bacterial clusters was significantly smaller than wildtype. High-magnification SD-IVM imaging demonstrated replication of wildtype staphylococci within KCs, whereas this intracellular growth phenotype was largely absent in ΔXV-infected mice (Fig. 2c). Importantly, this defect in intracellular growth was not explained by impaired bacterial catching, as both wildtype and ΔXV bacteria were taken up with equal efficiency by KCs during early stages of infection (Fig. 2d, Video 3). These findings indicate that TCS-dependent mechanisms are specifically required after bacterial internalization to support intracellular survival and replication. Accordingly, ΔXV-infected mice displayed significantly reduced bacterial burdens in the liver and kidneys 24 hours after infection compared with wildtype-infected animals (Fig. 2e) as well as reduced bacterial recovery from blood and peritoneal fluid (Supplemental Fig. 1b). Loss of TCS signalling also strongly attenuated staphylococcal virulence. All mice infected with a lethal dose of wildtype *S. aureus* succumbed to infection within 48 hours, whereas all mice infected with the ΔXV strain survived the duration of the experiment (Fig. 2f). Together, these findings demonstrate that non-essential TCS are critically important for intracellular replication within KCs and therefore mortality in systemic disease.

**Figure 2.**
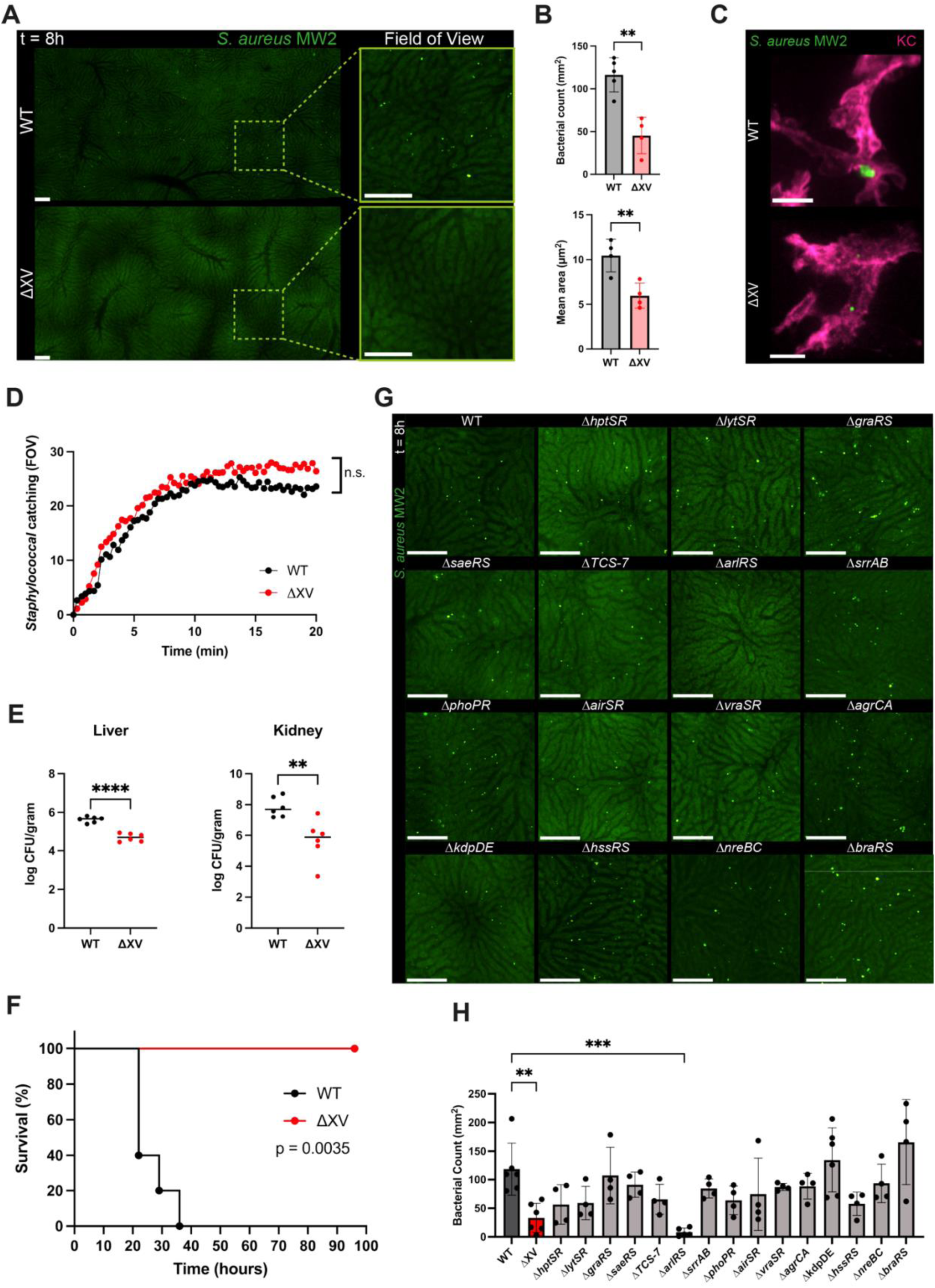
Identification of two-component systems required for intracellular replication in Kupffer Cells. (a) Representative large-stitched (left) and high-magnification (right) SD-IVM images of liver tissue 8 hours after intravenous infection with wildtype (WT; top panels) or ΔXV (bottom panels) *S. aureus* (5×10^7^ CFU) in C57Bl/6 mice. Scale bar 100 µm. (b) Quantification of bacterial microcolony number (top) and mean area (bottom) 8 hours after wildtype or ΔXV bloodstream infection (WT, n = 5 mice; ΔXV n = 4 mice). Unpaired t tests, mean +/-SD. ** p<0.01. (c) Representative SD-IVM 40× images of bacterial microcolonies in Kupffer cells 8 hours after wildtype (top) or ΔXV (bottom) infection. Kupffer cell (F4/80; magenta), *S. aureus* (GFP; green). Scale bar 25 µm. (d) Quantification of *S. aureus* capture following wildtype (black) or ΔXV (red) infection (WT, n = 3 mice; ΔXV, n = 4 mice). Welch’s t test. Data corresponds to **video S3**. (e) Bacterial burdens in liver and kidney 24 hours post intravenous infection with wildtype (black) or ΔXV (red) *S. aureus* (5×10^7^ CFU; n = 6 per group), unpaired t tests. Bars indicate median. (f) Survival following intravenous infection with wildtype (black) or ΔXV (red) *S. aureus* MW2 (1×10^8^ CFU; n = 5 mice per group). Kaplan-Meier survival analysis with Mantel-Cox test. (g) Representative SD-IVM images of liver 8 hours after wildtype, ΔXV, or individual single-TCS mutant strains. Scale bar 100 µm. (h) Quantification of bacterial microcolonies represented in (g). (WT, ΔXV, and Δ*kdpDE,* n = 6; Δ*arlRS* n = 5; all other strains, n = 4 mice). One-way ANOVA with multiple comparisons, mean +/-SD. ** p<0.01; *** p<0.001.

Previous studies have shown that individual TCSs frequently regulate bacterial adaptation to highly specific environmental stresses, including GraRS during acidic and antimicrobial peptide stress and SrrAB during hypoxia and nitrosative stress^22,26^. We therefore next asked whether a specific ‘single’ TCS is required for intracellular replication within Kupffer cells (KCs). Using SD-IVM, we analyzed intracellular microcolony formation of each individual TCS mutant strain at 8 hours post infection. Interestingly, deletion of the ArlRS TCS caused a near-complete defect in intracellular replication, whereas several other mutants, including Δ*hptSR*, Δ*lytSR*, Δ*TCS-7*, Δ*phoPR*, and Δ*hssRS*, showed only modest trends toward reduced microcolony formation (Fig. 2g,h). This was not due to altered GFP expression, as all strains expressed comparable levels of GFP driven by the constitutively active *sarA* promoter, as measured by flow cytometry (Supplemental Fig. 1c) These findings identify ArlRS as the dominant TCS required for *S. aureus* adaptation to the intracellular KC niche and suggest this is necessary for bacterial survival and replication.

### ArlRS TCS is essential for intracellular survival and replication

To further define the role of ArlRS during bloodstream infection, we directly compared intracellular bacterial replication and pathogenesis of wildtype, Δ*arlRS*, and chromosomally complemented Δ*arlRS* strains, each carrying the pCDM29 plasmid for constitutive eGFP expression. Large-stitched SD-IVM images of the liver 8 hours after infection with a sub-lethal dose of bacteria (5×10^7^ CFU) showed near elimination of Δ*arlRS* bacteria compared to wildtype or genetically complemented Δ*arlRS* strains (Fig. 3a). Quantification showed that intracellular colony formation was almost entirely abolished, whereas wildtype- and complemented strain-infected livers contained ∼50 microcolonies per mm^2^ (Fig. 3b). Furthermore, the few residual Δ*arlRS* microcolonies that were detected were significantly smaller averaging approximately one-third the size of wildtype colonies. High-resolution SD-IVM imaging further demonstrated the absence of intracellular staphylococcal clusters in Δ*arlRS*-infected KCs (Fig. 3c). This reduced intracellular burden was accompanied by a less inflammatory liver microenvironment, as Δ*arlRS-*infected mice recruited significantly fewer neutrophils compared to wildtype- and complemented strain-infected animals (Supplemental Fig. 1d). Again, the reduction in intracellular growth was not explained by impaired bacterial uptake, as all three strains were captured at equal rates by KCs (Fig. 3d, Video 4). In concordance with this, no differences in bacterial burden were observed in blood, liver, peritoneal fluid, spleen, or kidney 30 minutes after infection (Supplemental Fig. 1e), showing that in early stages of infection of there are no differences in organ localization.

**Figure 3.**
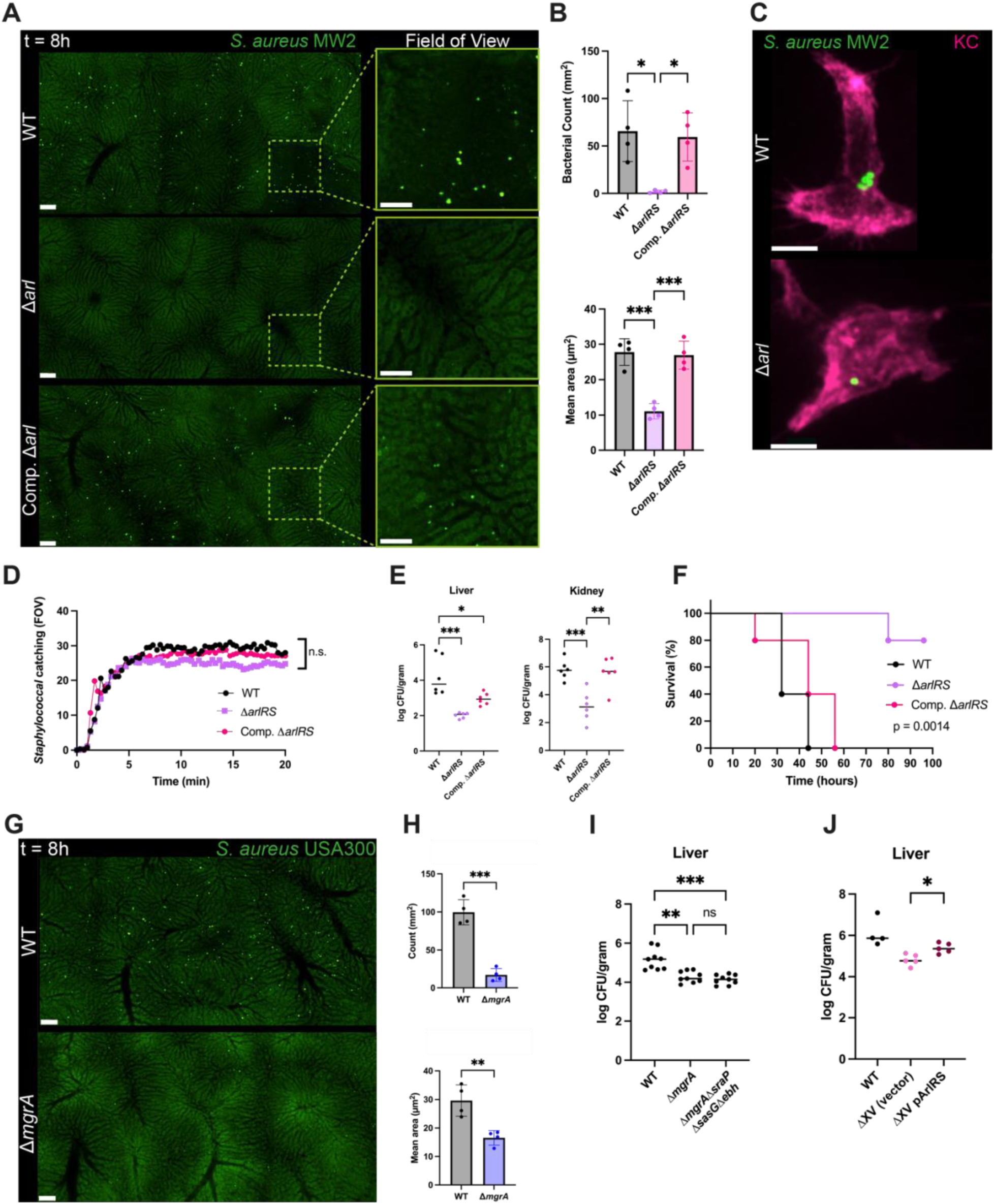
ArlRS is required for intracellular replication, virulence and pathogenesis during bloodstream infection. (a) Representative Large-field (left) and high-magnification (right) SD-IVM images of liver tissue 8 hours after intravenous infection with wildtype (WT; top panels), Δ*arlRS* (middle panels) or complemented Δ*arlRS* (bottom panels) *S. aureus* (5×10^7^ CFU). Scale bar, 100 µm. (b) Quantification of bacterial microcolony number (top) and mean microcolony area (bottom) 8 hours after infection. (n = 4 mice per group). One-way ANOVA, mean +/-SD. * p<0.05; *** p<0.001. (c) Representative 40× SD-IVM image of intracellular bacterial microcolonies in Kupffer cells following wildtype or Δ*arlRS* infection. Kupffer cell (F4/80; magenta), *S. aureus* (GFP; green). Scale bar 25 µm. (d) Quantification of *S. aureus* catching after infection with wildtype MW2 (black), Δ*arlRS* (violet), and complemented Δ*arlRS* (pink) strains (WT, n = 4 mice; ΔarlRS and complemented ΔarlRS, n = 5 mice). Brown-Forsythe and Welch ANOVA tests. Data correspond to **video S4.** (e) Bacterial burdens in liver and kidney 24 hours post intravenous infection with wildtype (black), Δ*arlRS* (violet), and complemented Δ*arlRS* (pink) *S. aureus* (5×10^7^ CFU; n = 6 mice per group), one-way ANOVA with multiple-comparison correction, bars indicate median. * p<0.05, ** p<0.01, *** p<0.001 (f) Survival following intravenous infection with wildtype, Δ*arlRS or* complemented Δ*arlRS S. aureus* (1×10^8^ CFU; n = 5 mice per group) Kaplan-Meier survival analysis with Mantel-Cox test. (g) Large SD-IVM (left) of liver tissue 8 hours after *S. aureus* USA300 LAC wildtype (WT; top) versus Δ*mgrA* (bottom) bloodstream infection (5×10^7^ CFU). Scale bar 100 µm. (h) Quantification of bacterial microcolony number (top) and mean area of bacterial microcolonies (bottom) following USA300 LAC wildtype (black) or Δ*mgrA* (blue) infection (n = 4 mice per group). Unpaired t tests, mean +/-SD. ** p<0.01, *** p<0.001. (i) Bacterial burden in liver 24 hours post intravenous infection with MW2 wildtype, Δ*mgrA*, or Δ*mgrA*Δ*sraP*Δ*sasG*Δ*ebh* (5×10^7^ CFU; WT, n = 9 mice). One-way ANOVA, mean +/-SD. ** p<0.01; *** p<0.001 (j) Bacterial burden in liver 24 hours post intravenous infection with MW2 wildtype, ΔXV carrying a control plasmid or ΔXV complemented with p*arlRS* (5 × 10^7^ CFU; WT, n = 4 mice; complemented strains, n = 5 mice). Brown-Forsythe and Welch ANOVA with multiple comparisons, median. * p<0.05.

The phenotypic difference between Δ*arlRS* and control strains persisted over time. At 24 hours after infection, Δ*arlRS-*infected mice exhibited a significantly reduced bacterial burden in both the liver and kidneys compared to wildtype-infected mice, whereas complementation with an *arlRS*-expressing plasmid mostly restored bacterial fitness (Fig. 3e). SD-IVM imaging of liver tissue at this later time point showed similar differences in staphylococcal expansion between the strains (Supplemental Fig. 1f). Consequently, loss of ArlRS profoundly attenuated virulence. Whereas all mice infected with a lethal inoculum of wildtype bacteria succumbed within four days, Δ*arlRS*-infected mice displayed markedly prolonged survival, a phenotype reversed by complementation (Fig. 3f). Collectively, these data identify ArlRS as a critical regulator of intracellular survival and replication within KCs and establish this TCS as a major determinant of *S. aureus* pathogenesis during bloodstream infection.

ArlRS has previously been linked to regulation of multiple virulence-associated processes, including autolysis, capsule production, adhesion, biofilm formation, and regulation of surface proteins^35–39^. More recently, transcriptomic studies demonstrated that ArlRS directly activates the global transcriptional regulator MgrA, thereby controlling a broad regulon involved in immune evasion and bacterial adaptation^40–42^. To determine whether MgrA contributes to intracellular replication within KCs, we analyzed a Δ*mgrA* mutant in the USA300 LAC background. Similar to Δ*arlRS* bacteria, Δ*mgrA* mutants displayed considerably reduced numbers and sizes of intracellular microcolonies following intravenous infection (Fig. 3g,h). These findings suggest that ArlRS-mediated adaptation to the intracellular KC environment is at least partially mediated through MgrA-dependent transcriptional regulation and further demonstrate that this phenotype is conserved across distinct *S. aureus* lineages. While MgrA is known to suppress the expression of several giant surface proteins, including Ebh, SasG, and SraP, which can mask adhesins and other surface-associated virulence factors^38^ and could therefore affect intracellular replication, deletion of all three genes in the *S. aureus* MW2 background failed to restore intracellular growth. (Fig. 3i). This indicates that these giant surface proteins are not responsible for the observed defect in intracellular replication. To determine whether ArlRS alone could restore intracellular fitness to the ΔXV mutant background, we complemented the *S. aureus* MW2 ΔXV strain with a plasmid expressing *arlRS*. Strikingly, restoration of ArlRS significantly increased bacterial burdens in the liver 24 hours after infection compared with ΔXV bacteria carrying an empty vector control (Fig. 3j). These findings indicate that ArlRS-dependent signaling is sufficient to substantially restore bacterial survival within KCs and suggest that the ArlRS regulon contains key adaptive pathways required for intracellular persistence during bloodstream infection.

The absence of intracellular microcolonies in Δ*arlRS*-infected mice could reflect either defective bacterial replication or impaired survival within the intraphagosomal environment. To distinguish between these possibilities, GFP-fluorescent strains were additionally labeled with N-Atto647 NHS ester, a fluorescent dye that binds covalently to the bacterial cell-wall and becomes diluted during bacterial division. Immunofluorescence analysis of infected liver sections demonstrated substantial dilution of the dye within wildtype intracellular microcolonies, indicating active bacterial replication within KCs (Fig. 4a, top panels). In contrast, Δ*arlRS* bacteria retained strong N-Atto647 labeling, indicating a profound defect in intracellular replication (Fig. 4a, bottom panels). Correspondingly, GFP-fluorescent bacteria, became increasingly abundant throughout wildtype microcolonies, whereas N-Atto647 signal remained present on Δ*arlRS* bacteria (Fig. 4b). Notably, some N-Atto647-associated fluorescent structures in Δ*arlRS*-infected livers lacked GFP expression entirely, indicating bacterial death. Super-resolution microscopy further enhanced visualization of cell-wall dye dilution and bacterial division events (Fig. 4c, Supplemental Fig. 2a). Wildtype bacteria displayed clear septal formation and incomplete N-Atto647 staining consistent with ongoing replication and daughter-cell formation. In contrast, Δ*arlRS* bacteria showed minimal dye dilution and frequently contained non-viable bacteria (Fig. 4c, Video 5). This approach allowed classification of bacteria into replicated (GFP^+^AF647^-^), dead (GFP^-^AF647^+^), and persistent non-replicating (GFP^+^AF647^+^) populations. Quantification revealed a dramatic reduction in total intracellular bacteria in Δ*arlRS*-infected livers and, importantly, a complete absence of replicated bacteria (Fig. 4d). Although both wildtype and Δ*arlRS* conditions contained persistent and dead bacterial populations, Δ*arlRS*-infected livers exhibited a higher proportion of dead bacteria and fewer persistent organisms. Collectively, these findings demonstrate that ArlRS is required not only for intracellular replication, but also for optimal bacterial survival within the KC intracellular niche.

**Figure 4.**
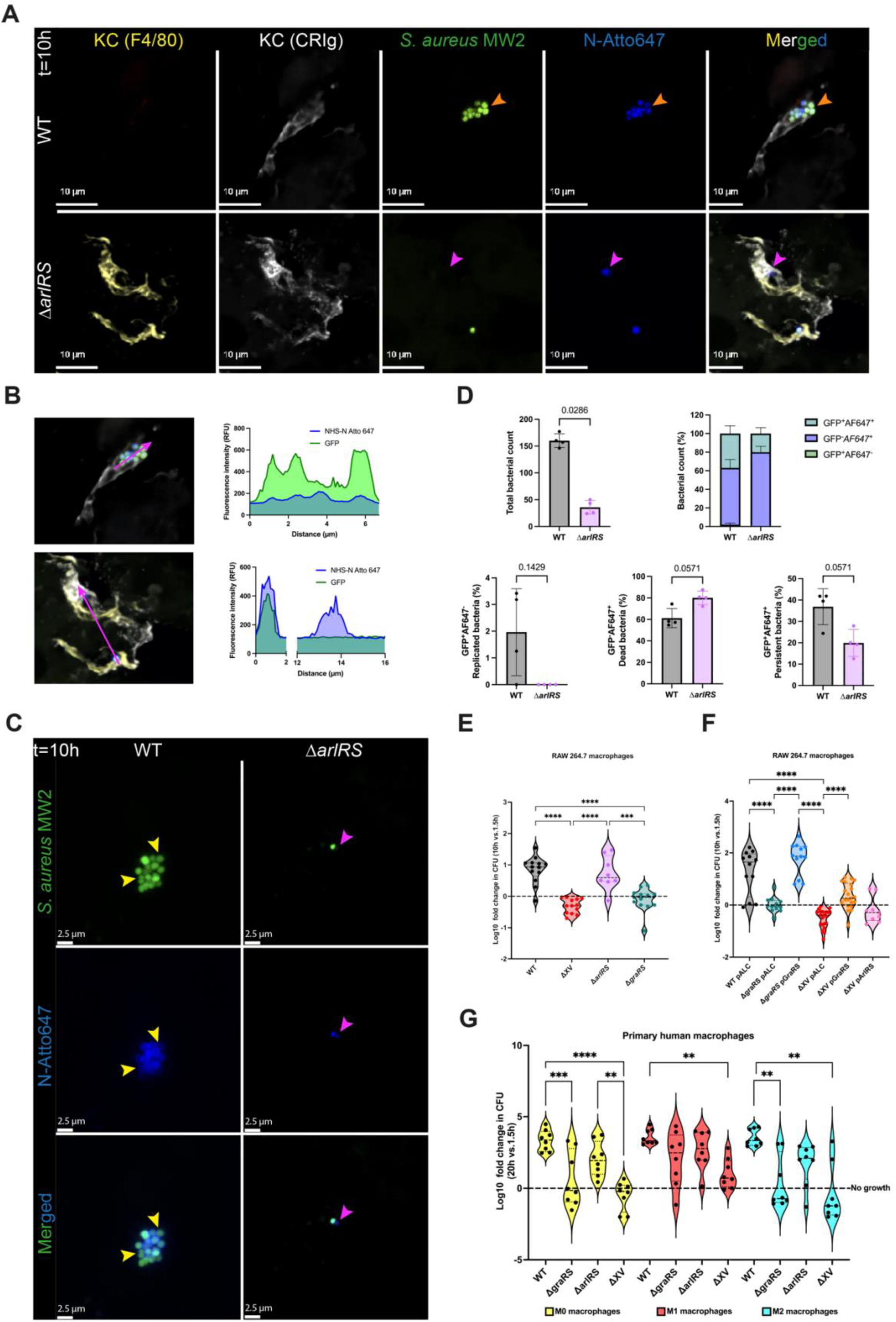
ArlRS is specifically required for bacterial replication within Kupffer cells. (a) Representative immunofluorescence image of liver cryosections from mice infected with wildtype (top) or Δ*arlRS*-infected (bottom) *S. aureus*. Bacterial replication was assessed by dual labeling of *S. aureus* with plasmid-encoded GFP and the covalently bound cell-wall dye NHS-N-Atto647, which is diluted upon bacterial division. Kupffer cells, F4/80 (yellow) and CRIg (white); *S. aureus*, GFP (green) and NHS-N-Atto647 (blue). Scale bar 10 µm. (b) Representative fluorescence intensity profile of GFP and NHS-N-Atto647 signals across intracellular wildtype (top) and Δ*arlRS* (bottom) bacteria inside KCs. Arrows indicate the regions used for analysis. (c) Representative STED image of whole-mount liver tissue from wildtype and Δ*arlRS*-infected mice. Dilution of the NHS-N-Atto647 signal is evident in replicating wildtype bacteria with septal formation (yellow arrows) but not in Δ*arlRS* bacteria. A dead bacterial cell can be observed in the Δ*arlRS* condition (magenta arrow). Scale bar, 2.5 µm. Data corresponds to **video S5**. (d) Quantification of bacterial populations in large liver cryosection images (1101.1 x 1101.1 µm) wildtype- and Δ*arlRS*-infected mice (n = 4 per group). Total bacterial counts per large image (top left) distribution of bacterial populations (top right) and frequencies of (replicated GFP^+^AF647^-^), dead (GFP^-^AF647^+^) and persistent (GFP^+^AF647^+^) bacteria (bottom panels). Kruskal-Wallis test for overview graph, mean +/-SD. Mann-Whitney test for other graphs, mean +/-SD. (e) Intracellular growth of wildtype MW2, ΔXV, Δ*arlRS*, and Δ*graRS S. aureus* in RAW264.7 macrophages (Δ*arlRS*, n = 8; all other groups, n = 13). One-way ANOVA with multiple comparisons. *** p < 0.001, **** p < 0.0001 (f) Intracellular growth of wildtype MW2, Δ*graRS*, complemented Δ*graRS*, ΔXV carrying a control plasmid, ΔXV complemented with pGraRS, and ΔXV complemented with pArlRS in RAW264.7 macrophages One-way ANOVA with multiple comparisons. **** p < 0.0001. (g) Intracellular growth of wildtype, Δ*graRS*, Δ*arlRS*, and ΔXV *S. aureus* in primary human M0, M1, and M2 macrophages (n = 8 donors). One-way ANOVA with multiple comparisons. ** p < 0.01, *** p < 0.001, **** p < 0.0001.

### ArlRS TCS importance is specific to the Kupffer cell intracellular environment

Previous work into intracellular staphylococcal replication identified the GraRS TCS as an important regulator of bacterial survival within RAW 264.7 macrophages^22^. To compare these findings with our *in vivo* KC imaging model we assessed intracellular growth of multiple TCS mutant strains by RAW 264.7 macrophages. As observed in KCs, the ΔXV strain displayed significantly impaired growth compared to wildtype bacteria. In agreement with Flannagan *et al.*, the Δ*graRS* showed considerably impaired intracellular growth^22^, surprisingly the Δ*arlRS* displayed no detectable effect on bacterial growth in this *in vitro* macrophage model (Fig. 4e). Complementation of Δ*graRS* with a plasmid expressing *graRS* fully restored intracellular replication, while overexpression of ΔXV with *graRS* partially rescued bacterial growth (Fig. 4f). In contrast, complementation of ΔXV with *arlRS* failed to improve intracellular replication within RAW264.7 macrophages.

To determine whether these observations extended to primary human macrophages, monocyte-derived macrophages were polarized into M0, M1 and M2 macrophage activation states before infection. Deletion of *graRS* significantly impaired intracellular growth in M0 and M2 macrophages, whereas deletion of *arlRS* had little effect (Fig. 4g). In contrast, neither *graRS* nor *arlRS* deletion affected intracellular growth in pro-inflammatory M1 macrophages. Nevertheless, the ΔXV mutant remained significantly attenuated in M1 cells, suggesting that an alternative TCS is required for intracellular adaptation these *in vitro* differentiated macrophages. Together, these findings demonstrate that the differentiation and activation state of macrophages dictates which bacterial sensory systems are required for intracellular survival and replication, highlighting the remarkable plasticity of *S. aureus* during adaptation to distinct intracellular niches. Notably, this contrasts sharply with our *in vivo* bloodstream infection model, in which ArlRS, but not GraRS, is essential for intracellular survival and growth within KCs (Fig. 2g,h).

### Loss of intracellular survival substantially enhances the efficacy of antibiotic therapy

Intracellularly residing bacteria are notoriously difficult to eradicate, as many antibiotics, including the last-resort glycopeptide vancomycin, exhibit poor penetration into KCs^4,43^. Consequently, intracellular *S. aureus* reservoirs remain largely unaffected by treatment, and following cessation of vancomycin therapy, surviving bacteria can resume replication, escape from KCs, and drive disseminated disease, including the formation of metastatic abscesses^4^. To test the impact of the intracellular bacterial compartment on pathogenesis, mice were infected i.v. with either wildtype or Δ*arlRS* bacteria and treated with 20 mg/kg i.v. vancomycin one-hour post-infection to eradicate any remaining extracellular bacteria. As expected, Δ*arlRS* infected animals all survived the infections over a period of 7 days, whereas most wildtype infected mice succumbed to infection irrespective of received antibiotics (Fig. 5a). The enhanced efficacy of vancomycin against the Δ*arlRS* mutant is not attributable to altered antibiotic susceptibility, as wildtype and Δ*arlRS S. aureus* display equivalent sensitivity to vancomycin with a MIC of ±1.5 µg/mL (supplemental Fig. 2c). Instead, loss of intracellular survival eliminates a protected bacterial reservoir that is refractory to antibiotic treatment. Bacterial burden in the kidney after 96 hours of infection showed that vancomycin treatment further reduced the bacterial presence of the Δ*arlRS* strain, and bacteria were fully absent in some mice (Fig. 5b), whereas the bacterial numbers of wildtype *S. aureus* were similar with or without vancomycin treatment. These data indicate that the intracellular KC reservoir specifically accounts for bacterial pathogenesis during systemic staphylococcal infection and that preventing the formation of this reservoir by removal of the ArlRS TCS significantly improved disease outcome.

**Figure 5.**
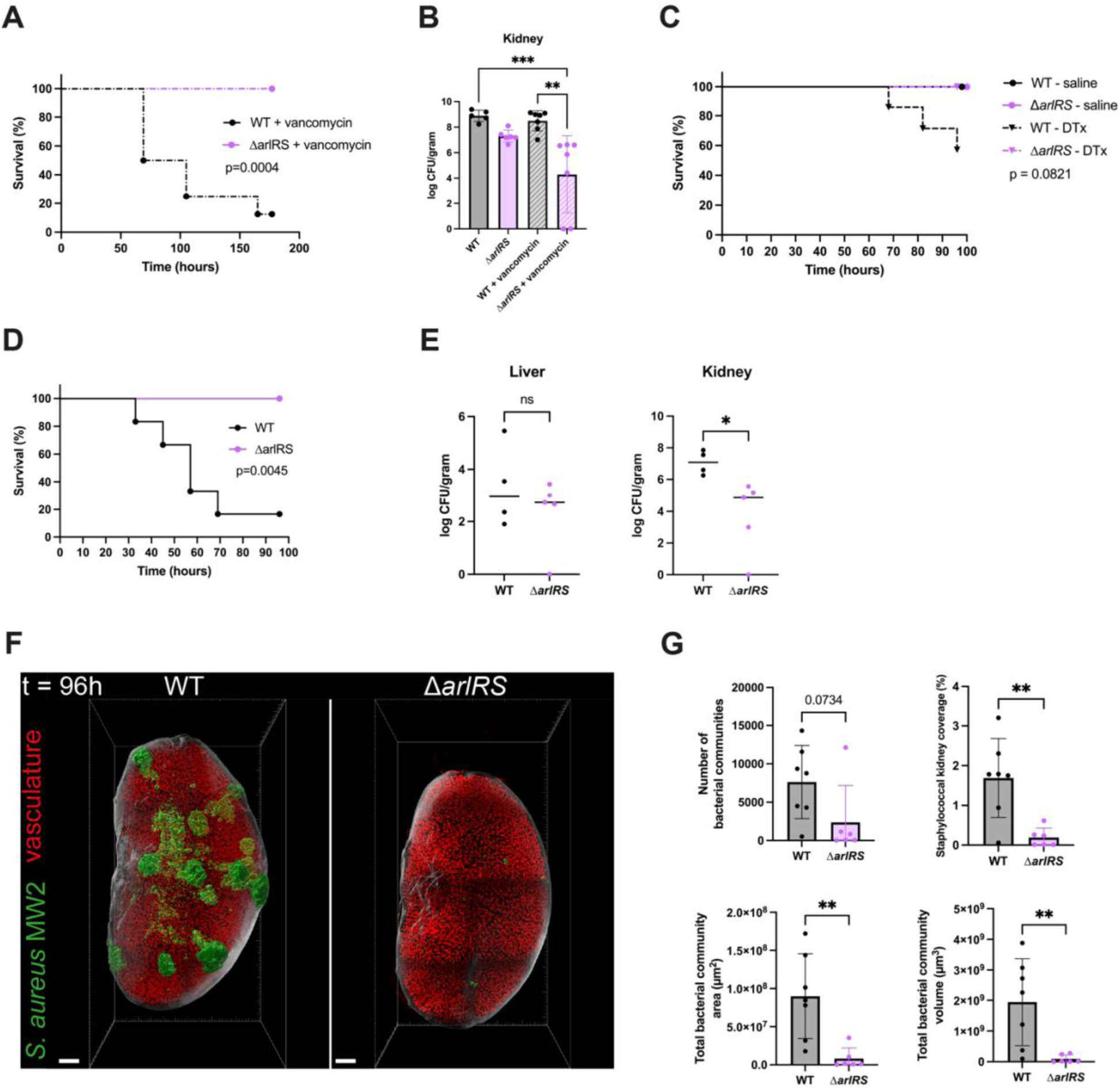
ArlRS is important in pathogenesis beyond the KC reservoir. (a) Survival of mice infected intravenously with wildtype or Δ*arlRS S. aureus* and treated with vancomycin (20 mg/kg, i.v.) 1 hour after infection (n = 8 mice per group). Kaplan-Meier survival analysis with Mantel-Cox test. (b) Kidney bacterial burden 96 hours after infection with wildtype or Δ*arlRS S. aureus* in saline-treated or vancomycin-treated mice. Vancomycin (20 mg/kg, intravenous) was administered 1 hour after infection (WT + saline, n = 5; Δ*arlRS* + saline, n = 6; all other groups, n = 7). Kruskal-Wallis test with multiple comparisons, mean +/-SD. ** p<0.01, *** p<0.001. (c) Survival of saline- or diphtheria toxin (DTx)-treated Clec4f-Cre-TdTomato^+/-^/iDTR^+/-^ mice following intravenous infection with wildtype or Δ*arlRS S. aureus* (5×10^7^ CFU). Solid lines indicate saline-treated mice, and dotted lines indicate DTx-treated mice (n = 2 for saline groups n = 6-7 DTx treated). Kaplan-Meier survival analysis with Mantel-Cox test. (d) Survival of clodronate liposome-treated mice infected intravenously with wildtype or Δ*arlRS S. aureus* at a dose of 5×10^6^ CFU (c; n = 6 mice per group). Kaplan-Meier survival analysis with Mantel-Cox test. (e) Liver and kidney bacterial burdens 24 hours after infection with wildtype or Δ*arlRS S. aureus* (5×10^6^ CFU) in clodronate liposome-treated mice (n = 4 mice per group). Bars indicate median values. * p<0.05. (f) Representative light-sheet images of cleared whole kidneys 96 hours after bloodstream infection with wildtype MW2 or Δ*arlRS S. aureus*. Bacteria (anti-LTA, green), vasculature (CD31, red), autofluorescence (white). Kidney tissue and bacterial abscesses were rendered as surfaces created in Imaris. Scale bar 700 µm. Data corresponds to **videos S6-S8**. (g) Quantification of total number of bacterial surfaces (top left), bacterial coverage of kidney tissue (top right), total abscess area (bottom left) and total abscess volume (bottom right) in wildtype- and Δ*arlRS*-infected kidneys (n = 6–7 mice per group) Mann-Whitney tests, mean +/-SD. ** p<0.01.

**ArlRS promotes systemic virulence through Kupffer cell-dependent and -independent mechanisms** The highly specific requirement for ArlRS within the KC niche suggests that the KC bottleneck itself is a major determinant for *S. aureus* virulence during bloodstream infection. To directly test this hypothesis, KCs were selectively depleted using Clec4f-Cre-tdTomato-iDTR mice. Using a lower infection dose (5×10^6^ CFU), all animals that reached the humane endpoint belonged to the diphtheria toxin-treated group infected with the wildtype strain (Fig. 5c), while all Δ*arlRS*-infected mice survived. This suggests that ArlRS also plays an important role in virulence beyond its role in establishing the KC reservoir. Consistent with this notion, broader depletion of resident macrophages and circulating monocytes using clodronate liposomes still revealed a reduction in virulence in the ArlRS mutant (Fig. 5d)^31,44^. To dive deeper into this phenotype, bacterial burdens were quantified 24 hours after infection in clodronate-treated mice. As expected, no difference in liver bacterial burden was observed between wildtype and Δ*arlRS*-infected animals, consistent with *S. aureus* not being able to establish a KC reservoir. In contrast bacterial burdens in the kidney were significantly reduced in Δ*arlRS*-infected animals in addition to lower counts in the spleen and bloodstream (Fig. 5e, supplemental Fig. 2b).

Given that the kidney is a major target organ for *S. aureus* bloodstream infection in mice, where bacteria escaping from KCs can seed the tissue and establish metastatic abscesses^4,45^, we next investigated whether loss of ArlRS affected renal colonization and abscess formation we next examined whether loss of ArlRS affected renal abscess formation. To achieve a comprehensive organ-wide assessment of infection burden, we employed whole-organ tissue clearing coupled with light-sheet microscopy, enabling 3D visualization and quantitative reconstruction of bacterial lesions throughout the entire kidney (Supplemental Fig. 2d). To our knowledge, this represents the first application of whole-kidney light-sheet imaging to assess the impact of a *S. aureus* virulence determinant on metastatic abscess formation following bloodstream infection. Unlike conventional histology, which samples only small tissue sections, this approach provides a quantitative assessment of the entire renal infectious burden. Using this methodology, we compared kidney pathology in mice infected with wildtype or Δ*arlRS* bacteria four days after intravenous challenge. Distinct anatomical regions of the kidney were readily identified from the XY plane of the three-dimensional light-sheet microscopy reconstruction (Supplemental Fig. 2e). Wildtype bacterial infection resulted in the formation of numerous bacterial clusters and abscesses distributed throughout the kidney (Fig. 5f, left panel, Video 6). In contrast, kidneys from Δ*arlRS*-infected mice showed a striking absence of bacterial abscesses (Fig. 5f, right panel, Video 7). Interestingly, this imaging approach showed the majority of bacterial abscess structures present in wildtype-infected kidneys to be located subcapsularly (Supplemental Fig. 2f). Some of these abscess structures penetrated deep into the tissue, into the renal medulla (Supplemental Fig. 2g, Video 8). Quantification of bacterial abscess structures revealed a trend towards a reduction of the total number of bacteria with Δ*arlRS* compared to wildtype infection (Fig 5g, top left). However, the absolute number of bacterial communities underestimated the magnitude of the phenotype, as larger clusters automatically resulted in fewer individual counts in our image analysis. Consistent with this, wildtype infection occupied a significantly greater proportion of kidney tissue, and individual abscess structures displayed markedly larger surface areas and volumes than those observed following Δ*arlRS* infection (Fig. 5g, right and bottom panels). Together, these findings demonstrate that ArlRS is essential for efficient renal colonization and abscess formation and establish that the contribution of ArlRS to *S. aureus* virulence extends beyond its role in promoting survival within the KC niche.

## Discussion

In this study, we identified the *S. aureus* ArlRS TCS as a critical regulator required for intracellular survival and growth within KCs, the specialized liver-resident macrophages that constitute an important reservoir during bloodstream infections. By using multiple complementary *in vivo* approaches, including intravital microscopy, bacterial genetics, KC depletion models, and whole-organ tissue clearing with light-sheet microscopy, we demonstrate that loss of ArlRS profoundly impairs bacterial adaptation to the KC intracellular environment. The attenuated phenotype was fully restored by genetic complementation, confirming that ArlRS directly mediates this virulence program. Failure to replicate within KCs prevented subsequent bacterial dissemination, resulting in improved mouse survival upon bloodstream infections, and almost complete absence of metastatic kidney abscesses. These findings establish ArlRS as a central determinant linking intracellular persistence within the liver to systemic disease progression.

A major finding of this study is that the bacterial sensory pathways required for intracellular adaptation are dictated by the host cell environment. While ArlRS was indispensable for survival and growth within KCs, it was dispensable in RAW264.7 macrophages and primary human macrophages. Conversely, GraRS, a well-established regulator of intracellular adaptation, was required in cultured M0 and M2 macrophages but not in KCs. Strikingly, neither GraRS nor ArlRS contributed to intracellular survival in inflammatory M1 macrophages, despite the observation that the ΔXV mutant remained attenuated, indicating that a distinct TCS likely mediates adaptation to this activation state. Together, these findings demonstrate that macrophage differentiation and tissue identity dictate which bacterial sensory pathways are required for intracellular adaptation. Given the remarkable heterogeneity of tissue-resident macrophages throughout the body, it is likely that *S. aureus* deploys distinct regulatory circuits to survive within different macrophage populations during infection.

KCs represent a highly specialized macrophage population whose identity is continuously maintained through interactions with liver sinusoidal endothelial cells, hepatocytes, stellate cells, circulating metabolites, and the gut microbiome^32,46^. These developmental and environmental cues shape lysosomal maturation, metabolism, antimicrobial mechanisms, and inflammatory responses, generating an intracellular environment that differs substantially from that encountered in monocyte-derived macrophages or immortalized cell lines. Our findings therefore highlight an important limitation of conventional macrophage infection models. Although these systems have been invaluable for identifying regulators such as GraRS, they do not fully recapitulate the specialized intracellular niche encountered during bloodstream infection. Consequently, virulence determinants that are essential *in vivo*, such as ArlRS, may remain undetected in simplified *in vitro* models.

Our data clearly demonstrated that ArlRS signaling is required for *S. aureus* adaptation to the intracellular KC environment. However, the downstream regulon responsible for the phenotype remains unknown. Previous transcriptomic studies have identified ArlRS-regulated genes, including the holin-like proteins lrgA and lrgB, capsule biosynthetic genes (*cap5A* and *cap5D)*, virulence factors such as *lukSF*, *lukAB*, and nuclease (*nuc*)^41,47^. The limited overlap between these studies suggest that the ArlRS regulon is highly dependent on environmental conditions, emphasizing the importance of defining ArlRS-dependent gene expression directly within infected KCs. Such analyses may uncover bacterial adaptations that are induced only within this specialized hepatic intracellular niche.

The host-derived signal that activates ArlRS within KCs remains unknown. Although ArlRS has been proposed to respond to magnesium^48^ and manganese starvation^49,50^, neither mechanism fully explains its activation during early stages of bloodstream infection. KCs rapidly expose internalized bacteria to a complex combination of nutritional limitation, oxidative stress, antimicrobial peptides, altered pH, host-derived lipids, and metabolic intermediates, any of which could contribute to ArlRS activation. It is also possible that ArlRS integrates multiple environmental cues rather than responding to a single stimulus. Defining both the environmental cues sensed by ArlRS and the downstream pathways it regulates will therefore be essential for understanding how *S. aureus* adapts to tissue-specific host environments.

Our study has several limitations. While we establish ArlRS as an essential regulator of intracellular adaptation within KCs, the downstream bacterial effectors and host-derived activating signals remain undefined. Likewise, although our findings demonstrate striking niche-specific regulation across macrophage populations, the bacterial sensory pathways that govern adaptation within other tissue-resident macrophages remain unknown. Future studies combining bacterial transcriptomics directly from infected KCs with genetic dissection of ArlRS-regulated pathways and characterization of the KC phagosomal environment will be necessary to define the molecular basis of this adaptation. Extending these studies to human liver models will also be important for determining the extent to which these mechanisms are conserved during human bloodstream infection.

Finally, our findings have important therapeutic implications. Intracellular persistence within KCs represents a critical bottleneck during *S. aureus* bacteremia and contributes to the limited efficacy of antibiotics with poor intracellular activity. Genetic disruption of *arlRS* alone markedly attenuated infection, and when combined with vancomycin treatment, infection became essentially avirulent. Although pharmacological inhibitors of ArlRS have previously been reported^51^, these compounds failed to reproduce the genetic phenotype in our *in vivo* model, likely due to limited intracellular penetration and insufficient target engagement within infected KCs. Consequently, additional studies aimed at developing inhibitors with improved intracellular delivery and validating their efficacy in physiologically relevant infection models will be required. Nevertheless, our findings suggest that targeting bacterial sensory pathways required for intracellular adaptation may represent an attractive anti-virulence strategy that complements existing antibiotic therapies while minimizing selective pressure for resistance.

Together, our work identifies the KC intracellular niche as a specialized environment that requires a dedicated bacterial adaptation program and demonstrates that the TCSs governing intracellular survival are determined by macrophage identity and activation state. These findings advance our understanding of how *S. aureus* senses tissue-specific host environments during bloodstream infection and underscore the importance of studying host-pathogen interactions in their native physiological context.

## Materials and Methods

### Mice

Adult mice, male and female (6-12 weeks old), were housed under specific pathogen-free conditions with a 12 h light/12 h dark cycle). C57Bl/6 mice were obtained from the Jackson laboratory and bred in-house. Clec4f-Cre-tdTomato-iDTR mice were generated by Clec4f-Cre-TdTomato (Jackson #33296) and ROSA26-iDTR (Jackson #7900) strains and double heterozygous offspring were used for experimentation. All procedures were approved by the University of Calgary Animal Care Committee (AC22-0049) and conducted in accordance with the Canadian Council on Animal Care.

### Antibodies and reagents

Anti-F4/80 (BM8), anti-Ly6G (1A8), anti-TIM-4 (RMT4-54) were obtained from BioLegend. Anti-CD31 PE (390), anti-CRIg (17C9) and anti-TIM-4 (21H12) were obtained from eBioscience. Anti-LTA^52^ was a kind gift from Brett Selman and conjugated with Alexa Fluor 555 protein labeling kit as per the manufacturer’s instructions. Anti-WTA antibody was kindly gifted by Dr. Suzan Rooijakkers.

### Spinning-disk intravital microscopy

For intravital microscopy, mice were anesthetized with ketamine/xylazine, catheterized via the tail vein for antibody administration, and prepared for liver imaging by laparotomy as previously described^53,54^. Intravital microscopy was performed using either an Olympus IX81 spinning-disk confocal microscope (Figs. 1–3) or a Nikon Ti2-E X1 spinning-disk microscope equipped with a Yokogawa CSU and ORCA Fusion BT sCMOS camera (Fig. 1 & 4). Images were acquired using 405, 488, 561, and 640 nm laser excitation and analyzed with NIS-Elements software (v5.42.04). For visualization purposes only, the 40× bacterial capture video was processed using AI denoising. Bacterial capture was quantified at 10× magnification in up to six randomly selected liver fields per mouse. Imaging was initiated 1 min before intravenous bacterial injection and continued for 25–30 min. High-resolution Z-stack images were acquired using a 60× water-immersion objective, whereas stitched images and collected at lower magnifications. Bacterial capture and burden were quantified from fluorescent signals using Volocity software (v7). Bacterial burden was assessed by measuring spot number and area, together with neutrophil counts, within a total imaging area of 2 mm². Bacterial replication was evaluated by comparing NHS-N Atto 647 and GFP fluorescence profiles across individual microcolonies using NIS-Elements software.

### Bacteria

*Staphylococcus aureus* strains used in this study are listed in **Table 1**. MW2 wildtype and mutant strains, including deletions in two-component regulatory systems and the 15-fold mutant (ΔXV), were provided by Dr. Iñigo Lasa. Unless otherwise indicated, these strains were transformed with plasmid pCM29 to enable constitutive expression of GFP, as previously described^6,55^. Select MW2 mutant strains carried either the pcn51 or parl plasmid, as specified.

**Table 1.** *Staphylococcus aureus* strains used in this study. MW2- and USA300-derived wildtype, mutant, complemented, and plasmid-expressing strains used in this study are listed, along with strain background and antibiotic resistance.

| Strain designation | Genetic description | Strain background | Plasmid | Antibiotic resistance |
| --- | --- | --- | --- | --- |
| WT | Wildtype | MW2 | pCM29 GFP | Chloramphenicol |
| ΔXV | 15-fold mutant | MW2 | pCM29 GFP | Chloramphenicol |
| ΔhptSR | HptSR deletion | MW2 | pCM29 GFP | Chloramphenicol |
| ΔlytSR | LytSR deletion | MW2 | pCM29 GFP | Chloramphenicol |
| ΔgraRS | GraRS deletion | MW2 | pCM29 GFP | Chloramphenicol |
| ΔsaeRS | SaeRS deletion | MW2 | pCM29 GFP | Chloramphenicol |
| ΔTCS-7 | TCS-7 deletion | MW2 | pCM29 GFP | Chloramphenicol |
| ΔarlRS | ArlRS deletion | MW2 | pCM29 GFP | Chloramphenicol |
| ΔsrrAB | SrrAB deletion | MW2 | pCM29 GFP | Chloramphenicol |
| ΔphoPR | PhoPR deletion | MW2 | pCM29 GFP | Chloramphenicol |
| **ΔairSR** | AirSR deletion | MW2 | pCM29 GFP | Chloramphenicol |
| **ΔvraSR** | VraSR deletion | MW2 | pCM29 GFP | Chloramphenicol |
| **ΔagrCA** | AgrCA deletion | MW2 | pCM29 GFP | Chloramphenicol |
| **ΔkdpDE** | KdpDE deletion | MW2 | pCM29 GFP | Chloramphenicol |
| **ΔhssRS** | HssRS deletion | MW2 | pCM29 GFP | Chloramphenicol |
| **ΔnreBC** | NreBC deletion | MW2 | pCM29 GFP | Chloramphenicol |
| **ΔbraRS** | BraRS deletion | MW2 | pCM29 GFP | Chloramphenicol |
| **ΔarlRS:pcn51** | ArlRS deletion with empty vector | MW2 | pcn51 | Erythromycin |
| **ΔarlRS:parl** | ArlRS deletion with *arlRS* complementation | MW2 | pArl | Erythromycin |
| **ΔXV:pcn51** | 15-fold mutant with empty vector | MW2 | pcn51 | Erythromycin |
| **ΔXV:parl** | 15-fold mutant with *arlRS* complementation | MW2 | pArl | Erythromycin |
| **WT** | Wildtype | MW2 | pCM29 GFP | Chloramphenicol |
| **ΔarlRS** | ArlRS deletion | MW2 | pCM29 GFP | Chloramphenicol |
| **comp ΔarlRS** | Complemented *arlRS* | MW2 | pCM29 GFP | Erythromycin;<br>Chloramphenicol |
| **WT** | Wildtype | USA300 | pCM29 GFP | Chloramphenicol |
| **ΔmgrA** | MgrA deletion | USA300 | pCM29 GFP | Chloramphenicol |
| **ΔmgrA** | MgrA deletion | MW2 | pCM29 GFP | Chloramphenicol |
| **ΔmgrA ΔsraP ΔsasG Δebh** | MgrA and large surface proteins deletion | MW2 | pCM29 GFP | Chloramphenicol;<br>Tetracycline |
| **WT** | Wildtype | MW2 |  |  |
| **ΔXV** | 15-fold mutant | MW2 |  |  |
| **ΔarlRS** | ArlRS deletion | MW2 |  |  |
| **ΔgraRS** | GraRS deletion | MW2 |  |  |
| **WT:pALC** | Wildtype with empty vector | MW2 | pALC | Chloramphenicol |
| **ΔgraRS:pALC** | GraRS deletion with empty vector | MW2 | pALC | Chloramphenicol |
| **ΔgraRS:pGraRS** | GraRS deletion with *graRS* complementation | MW2 | pGraRS | Chloramphenicol |
| **ΔXV:pALC** | 15-fold mutant with empty vector | MW2 | pALC | Chloramphenicol |
| **ΔXV:pGraRS** | 15-fold mutant with *graRS* complementation | MW2 | pGraRS | Chloramphenicol |
| **ΔXV:pArlRS** | 15-fold mutant with *arlRS* complementation | MW2 | pArlRS | Chloramphenicol |

Additional MW2 and USA300 strains, including Δ*mgrA*, complemented Δ*arlRS*, and associated derivatives, were kindly provided by Dr. Alexander Horswill.

### Cloning of *graRS* and *arlRS* in pALC2073

Complemented bacterial strains used for *in vitro* infection dynamics experiments were generated by cloning of *graRS* and *arlRS* into pALC2073 plasmids. The *graRS* and *arlRS* genes were PCR amplified from *S. aureus* MW2 using the following primer pairs for *graRS* (graR-KpnI: TATATA<u>GGTACC</u>GGAGAAAATATGAAATATACTAAATG with graS-SacI: TATATA<u>GAGCTC</u>CAAAACGCATGTTTAAAATGAC) and *arlRS* (arlR-KpnI: TATATT<u>GGTACC</u>CCTTATTAGAAAAAGTCGTAATATG with ArlS-SacI: TATATA<u>GAGCTC</u>GATCCTATGACTTTGATTGACG), respectively. The resulting amplicons were cloned into the KpnI/SacI site of *E. coli S. aureus* shuttle vector pALC2073 to generate the plasmids pGraRs and pArlRS that were maintained in *E. coli* DH5*a* with ampicillin at 100 µg/mL. Plasmids were verified by PCR and restriction analysis and then introduced into *S. aureus* RN4220 by electroporation using sucrose competent cells. Empty pALC2073 was electroporated and used as a vector control. Transformants were selected for in the presence of chloramphenicol at 10 µg/mL. Plasmids were isolated from *S. aureus* RN4220 and then electroporated into the desired *S. aureus* MW2 backgrounds using chloramphenicol selection.

### GFP expression analysis

Bacteria were grown until mid-exponential phase adiluted 10x in PBS 500 µL PB and analyzed by flow-cytometry (BD FACSCanto) Bacteria were gated based on forward and side scatter. GFP expression,was analyzed using FlowJo software (v. 10.10.1).

### Infections and *in vivo* treatments

For infection experiments, *S. aureus* strains were grown overnight in brain heart infusion medium (BD Bioscience) with shaking with chloramphenicol or erythromycin (10 µg/mL; BioShop) added where appropriate. Cultures were diluted and grown to mid-exponential phase, washed in sterile saline (Braun), and adjusted to an optical density an OD₆₆₀ of 1.0 (∼1×10⁹ CFU/mL). Final inocula were prepared by dilution immediately before infection. Mice were infected i.v. into the tail vein at 5 × 10^7^ CFU in 200 µL unless otherwise noted. Mice were monitored and sacrificed at the indicated time points. KC were depleted by DT treatment (10ng/g, i.p.) of Clec4f-Cre-iDTR mice whereas control mice received saline.

Global macrophage depletion was accomplished by injecting clodronate liposomes (200 µL i.v.) (LIPOSOMA BV) 48 hours before infection. Vancomycin 20 mg/kg (Sigma-Aldrich) was administered i.v. 1 hour after infection.

### Bacteriological analysis

Bacterial burdens in individual organs were quantified by colony-forming unit (CFU) analysis as previously described^56^. Blood was obtained by cardiac puncture and collected into 40 µL heparin (20 U/mL; BD Biosciences). Peritoneal lavage fluid was collected following intraperitoneal injection of 10 mL PBS and subsequently centrifuged for 10 min at 4,000 rpm; cell pellets were resuspended in 1 mL PBS. The spleen, liver, left kidney, and lungs were harvested and homogenized in 1 mL PBS using a tissue homogenizer (VWR 200). Blood and lavage volumes were recorded, and organ weights were determined before processing.

Samples were plated in serial 10-fold dilutions (starting at 1:10 or 1:100, depending on organ and time point) by spotting 30-µL aliquots in quadruplicate streaks onto blood agar plates. Plates were incubated overnight at 37°C in 5% CO₂. Colonies were enumerated from dilutions yielding 30–100 colonies, and bacterial loads were expressed as log₁₀ CFU per milliliter of fluid or per gram of tissue.

### Bacterial replication assessment

GFP-expressing bacteria were grown in a subculture as described previously and 1×10^9^ CFU bacteria were pelleted and labelled with 2 µL NHS ester N-Atto647 (Sigma-Aldrich) in 500 µL PBS for 30 minutes at 37ºC. Mice were infected for 8-10 h with 1×10^8^ CFU/200 µL. Liver samples were collected and 10 µm cryosections were prepared for immunofluorescence. GFP-expressing bacteria were grown in a subculture as described previously and 1 × 10^9^ CFU bacteria were pelleted and labelled with 2 µL NHS ester N-Atto647 (Sigma-Aldrich) in 500 µL PBS for 30 minutes at 37ºC. Bacteria were washed twice with PBS and diluted to a concentration of 5 × 10^8^ CFU to get an infectious inoculum of 1 × 10^8^ CFU/200 µL. 10 minutes prior to sample collection, mice were injected i.v. with fluorescent antibodies to label blood vessels. Mice were anesthetized by intraperitoneal injection of ketamine (200 µL/g; Vetoquinol) and xylazine hydrochloride (10 µL/g; Dechra). The abdomen was shaved and cut open to cannulate the interior vena cava. Once cannulated using polyethylene tubing (BTPE-90, Instech), the diaphragm was opened and the aorta was tied off using silk suture (5-0 USP, Look). Next, the portal vein was cut and 10 mL pre-warmed HBSS ([-] calcium chloride, [-] magnesium chloride, [-] magnesium sulfate, gibco) was pushed through using the KD legato 100 infuse single syringe pump (KD Scientific) to ensure constant and equal fluid pressure (4 mL/min; 30 mL BD plastic syringe). After HBSS, 10 mL of 10% formalin was flushed through the liver. The liver was removed and a small piece was cut off and put into a cassette (Simport Scientific Inc.) and fixed for 4 hours in 10% formalin. Samples were washed twice with PBS and then cryoprotected for 12-24 hours in 30% sucrose (Ward’s Science)/PBS. Samples were embedded into a disposable base mold (15x15x5 mm, Fisher Scientific) with a tissue embedding ring (Fisher Scientific) in Tissue Plus O.C.T. compound (Fisher Scientific) and stored at -80ºC until cryo-sectioning. One hour before sectioning, samples were moved to -20ºC and machine was cooled down to -20ºC. Samples were cut at 10 µm thickness using a cryostat machine and mounted on Superforst Plus glass microscope slides (Fisher Scientific). Slides were stored in -80ºC until immunofluorescence labelling. Slides were labelled with 100 µL 1:100 anti-F4/80 AF594 for at least 1 hour. Then, slides were washed 3 times with 0.1% Tween20 (Sigma-Aldrich)/PBS for 5 minutes rocking in the dark. Coverslip was mounted on the slides using ProLong Diamond Antifade Mountant (Life Technologies). Slides were labelled with 100 µL 1:100 anti-F4/80 AF594 for at least 1 hour. Sample images were collected with Z-stacks using a 93x/1.30 NA Glycerol lens on a Leica Stellaris 8 TauSTED microscope and Leica LAS X acquisition software. GFP and N-Atto647 were excited using the pulsed White Light Laser tuned to (488 and 640 nm) and N-Atto647 signal was depleted by the pulsed 775nm STED laser. The pixel size for sampling in STED resolution was set to 60 nm with a zoom factor of 4.15. 3D rendering and visualization were performed using Leica analysis software (Leica Application Suite X v. 5.3.1)

Whole-mount super resolution microscopy was performed with liver samples obtained from mice infected with dual-labelled bacteria after 8-10 hours of infection. 10 minutes prior to sample collection, Kupffer cells were labelled by i.v. administration of anti-F4/80 PE and anti-CRIg BV421. Mice were sacrificed and a piece of liver was taken up and fixed in 10% formalin for 30 minutes. The tissue was washed with PBS and immediately assessed with Leica Stellaris 8 TauSTED microscopy as described above.

### Macrophage tissue culture

Culture of RAW 264.7 murine macrophages (ATCC TIB-71™) and primary human macrophages was performed as previously described ^16^. In brief, macrophages were routinely maintained in RPMI 1640 with L-glutamine, 25 mM HEPES and sodium bicarbonate (Wisent) supplemented with 5% or 10% (v/v) FBS for RAW 264.7 and primary human macrophages, respectively. All macrophages were maintained in a humidified CO_2_ incubator at 37℃ and without antibiotics. For polarization of macrophages into M1, M0 and M2 states the following cytokines were used: For M1, GM-CSF (20 ng/mL), human IFN-ɣ (10 ng/mL, Peprotech) and LPS (250 ng/mL, Sigma Aldrich) was used; For M0, only M-CSF (20 ng/mL) was used; For M2 M-CSF (20 ng/mL) and IL-4 (15 ng/mL) was used. Whole human blood for primary macrophage culture was drawn from healthy adult volunteers upon verbal consent in accordance with protocols approved by the University of Western Ontario’s Research Ethics Board, London, Ontario, Canada.

### Macrophage phagocytosis assays

For RAW 264.7 macrophage infection phagocytes were split by scraping and seeded into 12-well tissue culture-treated dishes as previously described ^16^. In brief, macrophages were infected at an MOI of 30 for RAW 264.7 cells or 10 for primary human macrophages. Tissue culture plates with macrophages and bacteria were centrifuged at 277 x g for 2 min then incubated for 30 min at 37℃. Next, the medium was aspirated and replaced with SF-RPMI containing gentamicin (100 µg/mL) and incubated for 1 hour. Samples were tested for bacterial enumeration at 1.5 hours or 10 hours postinfection for RAW cells or 20h post-infection for human macrophages. Once the desired time point was reached, cells were lysed in 500 µL of 0.1% (v/v) triton x-100 in PBS. These lysates were serially diluted and plated onto TSA and incubated O/N at 37℃ to enable bacterial enumeration. The ability of macrophages to restrict bacteria growth was expressed as a log10 fold change in CFU where the bacterial counts for each replicate at 10 hpi or 20 hpi were divided by the counts at 1.5 hpi and the resulting value log10 transformed.

### Kidney abscess visualization

Mice were infected as previously described with 5×10^7^ CFU bacteria for 96 hours. 10 minutes prior to sample collection, mice were injected i.v. with anti-CD31 AF647 to label blood vessels. Using an electric syringe pump as described in the supplemental methods, whole-body perfusion was performed by inserting a 20g syringe (BD Bioscience) into the left ventricle and making a cut in the right atrium. The body was perfused using 20 mL PBS and subsequently with 20mL 4% PFA (Thermo Scientific)/PBS. Tissue dissection was performed to remove both kidneys, which were prepared for lightsheet imaging using the iDISCO tissue clearing protocol^57^. Kidney tissues were fixed overnight in 4% PFA/PBS at 4ºC while shaking. Samples were dehydrated with methanol (Fisher Scientific)/H2O series: 25%, 50%, 75%, 100%, 100%; 1 hour each at RT while shaking. Samples were cooled to 4ºC and transferred to TraceClean 20 mL glass vials (VWR). Samples were incubated overnight in 66% DCM (Sigma-Aldrich)/33% methanol at RT while shaking. Samples were washed twice with 100% methanol for 30 minbuntes at RT while shaking, then chilled to 4ºC. Samples were bleached in chilled, fresh 5%H2O2 (Sigma-Aldrich) in methanol overnight at 4ºC. Samples were rehydrated with methanol/H2O series: 75%, 50%, 25%, PBS; 1h each at RT while shaking. Samples were washed twice with PTx.2 (100 mL PBS 10X, 2 mL TritonX-100 (Sigma-Aldrich), total 1L volume) at RT. Samples were incubated with permeabilization solution (400 mL PTx.2, 11.5 g glycine (VWR), 100 mL DMSO (Sigma-Aldrich)) for 18-24 hours at 37ºC after which they were blocked in blocking solution (42 mL PTx.2, 3 mL donkey serum (Sigma-Aldrich), 5 mL DMSO) for 24-48h at 37ºC. Samples were incubated with 1:500 anti-LTA Alexa Fluor 555 to stain bacteria in PTwH (100 mL PBS 10X, 2 mL Tween20 (Sigma-Aldrich), 1 mL 10 mg/mL heparin stock solution (Sigma-Aldrich))/5% DMSO/3% donkey serum at 37ºC for 24-48 hours. Samples were washed 4-5x with PTwH over a period of 24 hours. Samples were dehydrated with methanol (Fisher Scientific)/H2O series: 25%, 50%, 75%, 100%; 1 hour each at RT while shaking, then another 100% methanol overnight at RT while shaking. Samples were incubated in 66% DCM/33% methanol at RT for 3 hours shaking. Samples were incubated in 100% DCM 15 minutes twice while shaking to wash away methanol. Samples were incubated in DiBenzyl Ether (Sigma-Aldrich) until imaging. On the day of imaging, kidney samples were transferred into Ethyl cinnamate, 98+% (Thermo Fisher Scientific). Kidneys were stained with 1:500 anti-LTA Alexa Fluor 555 to label bacteria. Cleared kidneys were imaged using a LaVision BioTec UltraMicroscope II equipped with a 4× objective, multiple laser lines (405–785 nm), and an Andor Zyla 4.2 Plus sCMOS camera. Images were acquired with ImSpector software, stitched, and analyzed in Imaris (v11.0.1). Kidney volume, abscess number, size, and total abscess volume were quantified.

### Statistical analysis

Statistical analysis was performed using GraphPad Prism (v11) software. For comparison between two groups, the unpaired Student’s t-test was used. For comparisons between two groups over multiple time points, multiple Mann-Whitney tests were performed according to the Holm-Šídák method. Comparisons performed over three groups were tested with a Kruskal-Wallis test with multiple comparisons. The Kaplan-Meier test was performed together with a Log-rank (Mantel-Cox) test for survival curve analysis.

## Acknowledgements

We thank Trecia Nussbaumer and Luisa Samaniego Barrón for the breeding of mice. We thank Dr. Karen Poon at the Flow cytometry core facility at the University of Calgary for her help with flow cytometry. Many thanks to Dr. Ysbrand Nusse for helping with cryosectioning. We acknowledge the Hotchkiss Brain Institute Advanced Microscopy Platform and the Cumming School of Medicine for support and use of LaVision BioTec UltraMicroscope II, as well as the service provided by staff scientist David Elliott for their help with microscopy setup and image analysis and visualisation. Lastly, our gratitude to Lianne Dale and Gregory Plummer from the Leica support team for their help with STED super resolution microscopy.

B.G.J. Surewaard received funding through University of Calgary start-up funds, Canada Research Chairs program, NSERC Discovery Grant and Snyder Institute catalyst Grants. J.W. Hommes is partially funded by the Achievers in Medicine Doctoral Scholarship of the University of Calgary.

The authors declare no competing financial interests.

Author contributions: J.W. Hommes and B.G.J. Surewaard designed the study and wrote the manuscript. J.W. Hommes, L.M. Orthner, F. van der Wal, R.S. Flannagan, R. Shim, L. Samaniego Barrón, and M. Dao performed experiments. J.W. Hommes performed data analysis and visualization. I. Lasa, and A.R. Horswill provided bacterial strains used for experiments. D.E. Heinrichs and B.G.J. Surewaard acquired funding. All authors provided edits for the manuscript.

## Notes

### Competing Interest Statement

The authors have declared no competing interest.

