## Supplemental Figures for "A niche-specific role for ArlRS enables *Staphylococcus aureus* growth within Kupffer cells"

### *Staphylococcus aureus* ArlRS two-component system is required for intracellular survival and growth within Kupffer cells

Josefien W. Hommes[1], Lindsey M. Orthner[1], Fardau van der Wal[1], Ronald S. Flannagan[2], Raymond Shim[1], Luisa Samaniego Barrón[1], Mai Dao[1], Iñigo Lasa[3], Alexander R. Horswill[4], David E. Heinrichs[2], Bas G. J. Surewaard[1]*

[1] = Department of Microbiology, Immunology, and Infectious Disease. Snyder Institute for Chronic Diseases, University of Calgary, Calgary, AB, Canada.

[2] = Department of Microbiology and Immunology, The University of Western Ontario, London, Ontario, Canada

[3] = Laboratory of Microbial Pathogenesis, Navarrabiomed-Hospital Universitario de Navarra (HUN)-Universidad Pública de Navarra (UPNA), IDISNA, Pamplona, Navarre, Spain

[4] = Department of Immunology and Microbiology, University of Colorado School of Medicine, Aurora, Colorado, USA

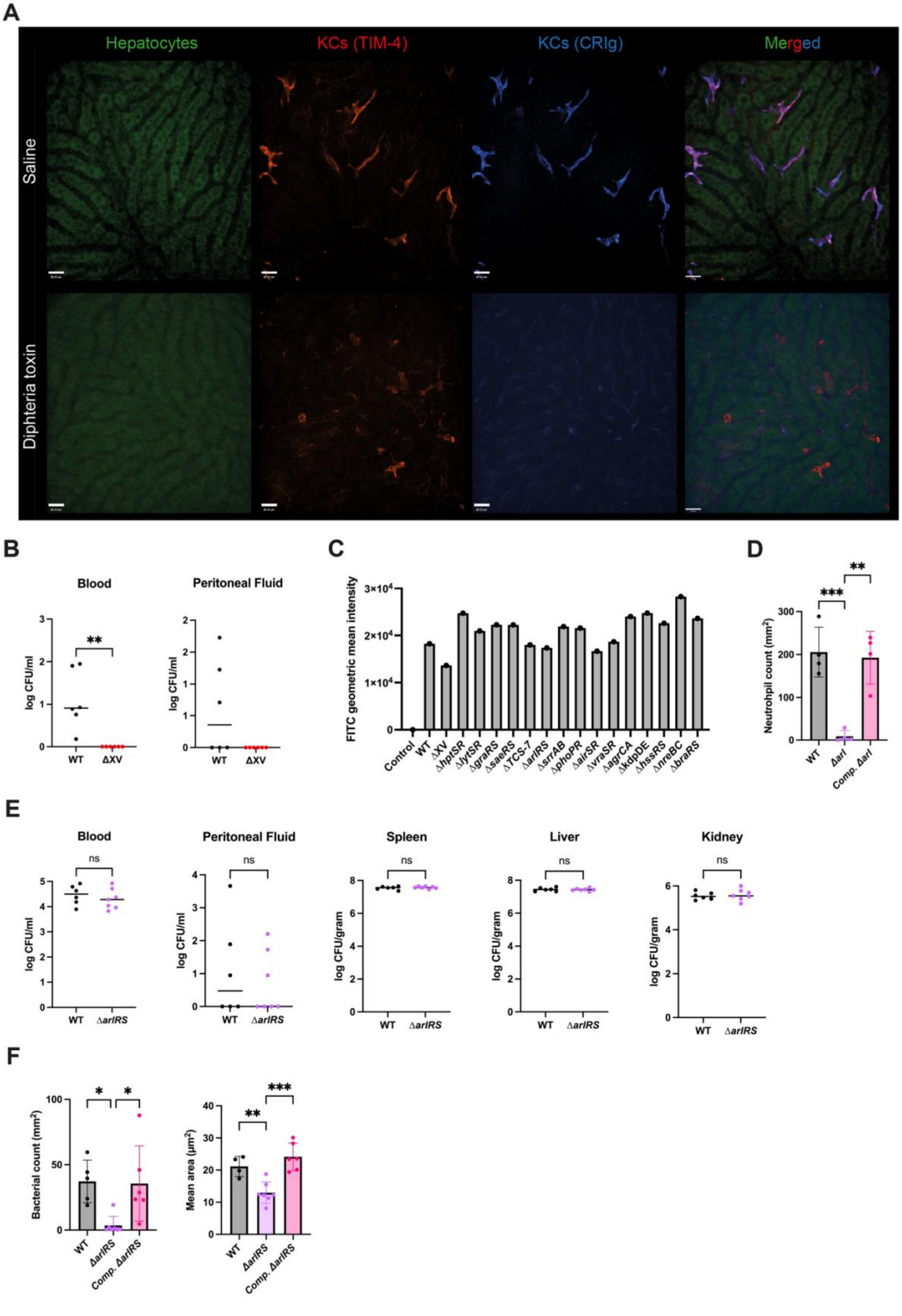

A
Hepatocytes
KCs (TIM-4)
KCs (CRIg)
Merged
Saline
Diphteria toxin
B
Blood
Peritoneal Fluid
log CFU/ml
WT
ΔXV
**
C
FITC geometric mean intensity
3×10⁴
2×10⁴
1×10⁴
0
Control
WT
ΔXV
ΔhptSR
ΔlytSR
ΔgraRS
ΔsaeRS
ΔTCS-7
ΔarlRS
ΔsrrAB
ΔphoPR
ΔairSR
ΔvraSR
ΔagrCA
ΔkdpDE
ΔhssRS
ΔnreBC
ΔbraRS
D
Neutrohpil count (mm²)
WT
Δarl
Comp. Δarl
***
**
E
Blood
Peritoneal Fluid
Spleen
Liver
Kidney
ns
log CFU/ml
log CFU/gram
WT
ΔarlRS
F
Bacterial count (mm²)
Mean area (µm²)
WT
ΔarlRS
Comp. ΔarlRS
*
**
***

**Supplemental figure 1. Two-component system*s* are required for intracellular bacterial growth of *Staphylococcus aureus*.** (a) SD-IVM images of liver tissue from Clec4f-Cre-TdTomato$^{+/-}$/iDTR$^{+/-}$ mice 12 hours after treatment with saline (top) or diphtheria toxin (bottom) Hepatocytes (autofluorescence; green), Kupffer cells (TIM-4; red and CRIg; blue). Scale bar 25 µm. (b) Bacterial burdens in blood and peritoneal fluid 24 hours post intravenous infection with wildtype or ΔXV *S. aureus* ($5\times10^7$ CFU; n = 6 mice per group), unpaired t tests; median. (c) GFP expression of wildtype, ΔXV, and individual TCS mutant strains represented as the geometric mean intensity. (d) Quantification of recruited neutrophils 8 hours after bloodstream infection with wildtype, Δ*arlRS,* and complemented Δ*arlRS S. aureus* (n = 4 mice per group). One-way ANOVA, mean +/- SD. ** p<0.01; *** p<0.001. (e) Bacterial burdens in blood, peritoneal fluid, liver, and kidney 30 minutes post intravenous infection with wildtype or Δ*arlRS S. aureus* ($5\text{x}10^7$ CFU; n = 6 mice per group), one-way ANOVA with multiple comparisons correction, Bars indicate median values. (f) Quantification of number of bacterial microcolonies (left) and mean microcolonies area (right) 24 hours after bloodstream infection with wildtype, Δ*arlRS*, and complemented Δ*arlRS S. aureus* strains (n = 4 mice per group). One-way ANOVA, mean +/- SD. * p<0.05; ** p<0.01; *** p<0.001.

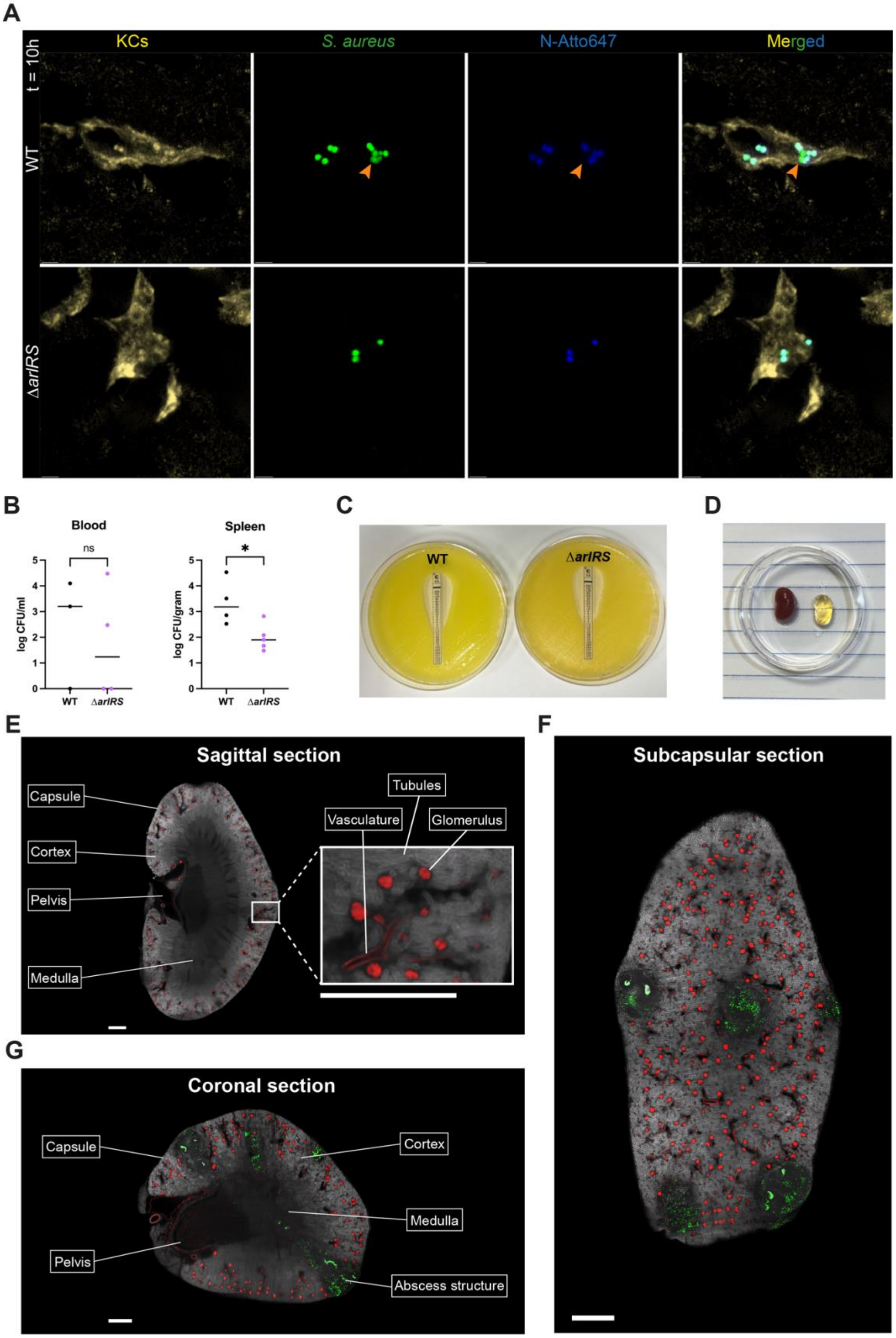
A
KCs
S. aureus
N-Atto647
Merged
t = 10h
WT
ΔarlRS
B
Blood
Spleen
ns
*
log CFU/ml
log CFU/gram
WT
ΔarlRS
C
WT
ΔarlRS
D
E
Sagittal section
Capsule
Cortex
Pelvis
Medulla
Tubules
Vasculature
Glomerulus
F
Subcapsular section
G
Coronal section
Capsule
Cortex
Medulla
Pelvis
Abscess structure

**Supplemental figure 2. ArlRS is required for bacterial intracellular growth.** (a) STED super-resolution image of liver cryosections of intravenously infected with WT (top) or ΔarlRS *S. aureus* (bottom). Bacterial replication was assessed using the cell wall dye N-Atto647-NHS ester, which covalently labels the bacterial cell wall. Because the dye is diluted with each round of bacterial division, loss of signal indicates bacterial replication (orange arrows). KCs (F4/80, yellow; CRIg, white), *S. aureus* (GFP, green), NHS-ester (N-Atto647, blue). (b) Bacterial burden in blood and spleen tissue 24 hours post intravenous infection with *S. aureus* wildtype (black) versus Δ*arlRS* (violet) ($5\times10^6$ CFU) in mice treated with clodronate liposomes, (n = 4 per group), Mann-Whitney tests, median. * $p<0.05$. (c) Vancomycin susceptibility of both *S. aureus* MW2 wildtype and Δ*arlRS* using a MIC test strip. (d) Representative photo of normal healthy kidney (left) and kidney after tissue clearing using the iDISCO protocol (right). (e) Representative light-sheet microscopy image of sagittal plane of a kidney after 96 hours of infection with *S. aureus* MW2 wildtype. Different kidney compartments were labelled based on the tissue structure (autofluorescence, white). Vasculature was labelled using CD31 AF647 (red). Scale bars 700 µm. (f) Representative light-sheet microscopy image of subcapsular plane of a kidney after 96 hours of infection with *S. aureus* MW2 wildtype. Large bacterial abscess structures were visualized (anti-LTA, green). Tissue was visualized based on autofluorescence (white) and vasculature was labelled using CD31 AF647 (red). Scale bar 700 µm. (g) Representative light-sheet microscopy image of coronal plane of a kidney after 96 hours of infection with *S. aureus* MW2 wildtype. Different kidney compartments were labelled based on the tissue structure (autofluorescence, white). Vasculature was labelled using CD31 AF647 (red). Bacterial abscess structures were visualized using anti-LTA (green). Scale bar 500 µm.

**Video 1. Staphylococcal catching by Kupffer cells.**

Representative 40× SD-IVM video of a murine liver over 20 minutes. At 1 minute, $5\times10^7$ CFU *S. aureus* MW2 wildtype-GFP (green) was injected intravenously. Kupffer cells (TIM-4 BV421, magenta) situated in the liver sinusoids (CD31 AF647; white) rapidly captured circulating bacteria.

**Video 2. Staphylococcal catching is impaired when KCs are absent.**

Representative 10× SD-IVM video of murine liver tissue over 20 minutes. At 2 minutes, $5\times10^7$ CFU *S. aureus* MW2 wildtype-GFP (green) was injected intravenously into Clec4f-Cre-TdTomato$^{+/-}$/iDTR$^{+/-}$ mice. Saline-treated control animals (first video) rapidly captured circulating bacteria within the liver. In contrast, diphtheria toxin-treated animals depleted of Kupffer cells (second video) showed reduced bacterial capture. Kupffer cells were visualized using TIM-4 PE (red) and CRIg AF647 (blue).

**Video 3. *S. aureus* MW2 wildtype and ΔXV bacteria are captured by KCs with equal efficiency.**

Representative 10× SD-IVM video of murine liver tissue over 25 minutes. At 1 minute, $5\times10^7$ CFU *S. aureus* MW2 wildtype (first video) or ΔXV (second video) were injected intravenously into C57Bl/6 mice. Bacteria were rapidly captured by liver Kupffer cells at equal efficiency. Kupffer cells (F4/80 AF750, purple) *S. aureus* (GFP, green).

**Video 4. *S. aureus* MW2 wildtype, ΔarlRS, and complemented ΔarlRS bacteria are captured by Kupffer Cells with equal efficiency.**

Representative 10× SD-IVM video of murine liver tissue over 20 minutes. At 1 minute, $5\times10^7$ CFU *S. aureus* MW2 wildtype (first video), Δ*arlRS* (second video), or complemented Δ*arlRS* (third video) was injected intravenously into C57Bl/6 mice. Bacteria were rapidly captured by liver Kupffer cells at similar rates for all strains. Kupffer cells (F4/80 AF750, purple) *S. aureus* MW2 (GFP, green).

**Video 5. *S. aureus* MW2 ArlRS TCS is required for intracellular bacterial growth.**

Representative Leica Stellaris 8 STED super resolution videos of bacterial microcolonies in whole mount liver tissue 10 hours post bloodstream infection with *S.aureus* MW2 wildtype (first video) formed large grapelike intracellular microcolonies. Dilution of the cell wall dye N-Atto647 NHS ester (AF647 STED, purple) among GFP + daughter bacteria indicated active bacterial replication. In contrast, *S. aureus* Δa*rlRS* (second video) bacteria failed to form microcolonies and retained N-Atto647 NHS ester labeling, indicating an absence of bacterial replication. Bacteria lacking GFP signal were also observed, consistent with bacterial cell death. Three-dimensional rendering of STED signal using Leica Application Suite X.

**Video 6. *S. aureus* MW2 wildtype grows large abscesses in murine kidney.**

Representative light-sheet video of cleared kidney tissue 96 hours post intravenous infection with *S. aureus* MW2-wildtype. 3D renderings of kidney autofluorescence (488 nm) and bacterial abscesses (anti-GFP AF555) were generated using Imaris software. Numerous Large bacterial abscesses were observed throughout the kidney tissue following WT *S. aureus* infection. Vasculature was visualized using CD31 AF647 (red).

**Video 7. *S. aureus* MW2 ΔarlRS kidney abscess formation is impaired.**

Representative light-sheet video of cleared kidney tissue 96 hours after intravenous infection with *S. aureus* MW2 Δ*arlRS*-GFP. 3D renderings of kidney autofluorescence (488 nm) and bacterial abscesses (anti-GFP AF555) were generated using Imaris software. Few bacteria were detected within the kidney tissue, and the bacterial clusters that were present were markedly smaller and failed to form dense abscess structures than those observed following wildtype infection. Vasculature was visualized using CD31 AF647 (red).

**Video 8. Bacterial distribution within the kidney.**

Representative light-sheet video of cleared kidney tissue 96 hours post intravenous infection with *S. aureus* MW2-wildtype. 3D renderings of kidney autofluorescence (488 nm) and bacterial abscesses (anti-GFP

AF555) were generated using Imaris software. Vasculature was visualized using CD31 AF647 (red). Slice mode was used to show individual kidney planes both in XY and XZ plane. Large abscess structures can be observed just below kidney capsule. These structures penetrate deeper into the kidney medullar structure.
